# Male-specific features of *C. elegans* neuronal aging

**DOI:** 10.1101/2023.12.18.572229

**Authors:** Yifei Weng, Coleen T. Murphy

**Affiliations:** Department of Molecular Biology University, Princeton, NJ 08544; LSI Genomics, Princeton University, Princeton, NJ 08544

## Abstract

Aging is a complex biological process, with sexually dimorphic aspects. For example, men and women differ in their vulnerabilities in cognitive decline, suggesting biological sex may contribute to the heterogeneous nature of aging. Although we know a great deal about the cognitive aging of hermaphrodites of the model system *C. elegans,* less is known about cognitive decline in males. Through behavioral analyses, we found that the cognitive aging process has both sex-shared and sex-dimorphic characteristics. Through neuron-specific sequencing, we identified neuronal age-associated sex-differential targets. In addition to sex-shared neuronal aging genes, males differentially downregulate mitochondrial metabolic genes and upregulate GPCR genes with age. In addition, the X chromosome exhibits increased gene expression in hermaphrodites and altered dosage compensation complex expression with age, indicating possible X-chromosomal dysregulation that contributes to sexual dimorphism in cognitive aging. Finally, we found that the sex-differentially expressed gene *hrg-7*, which encodes an aspartic-type endopeptidase, regulates male behavior during cognitive aging but does not affect hermaphrodites’ behaviors. Overall, these results suggest that males and hermaphrodites exhibit different age-related neuronal changes. This study will strengthen our understanding of sex-specific vulnerability and resilience and help identify new pathways to target with novel treatments that could benefit both sexes.

## Introduction

Cognitive decline with age is a growing health issue as human life expectancy continues to increase (World Health Organization, 2021). Sexual dimorphism contributes to the heterogeneity of this process. In humans, women generally outlive men (Thornton, 2019), and various biological markers of aging demonstrate sex differences (Hägg and Jylhävä, 2021), and are regulated both by sex chromosomes and sex hormones (Gentilini et al., 2012; Davis et al., 2020). The nervous system is an important target of aging, not only affecting lifespan but also affecting the quality of life, and brain aging also exhibits sex-specific features. Women exhibit slower biological brain aging (Horvath et al., 2016; Goyal et al., 2019), less structural damage (O’Dwyer et al., 2012), and decreased cognitive ability decline (Jack et al., 2015; Casaletto et al., 2019). In addition, men and women show different frailty toward neurodegenerative diseases (Yang et al., 2022). However, a comprehensive molecular characterization is required to compare the neuronal aging trajectory between sexes.

*C. elegans* lives for two to three weeks, and displays many quantifiable aging phenotypes during this time (Garigan et al., 2002; Herndon et al., 2002; Huang et al., 2004; Murakami and Murakami, 2005; Hughes et al., 2007). *C. elegans* populations are primarily hermaphroditic, but about one in 500 *C. elegans* is a male under unstressed conditions (Hodgkin et al., 1979). Both sexes have invariant, stereotyped nervous systems that have been fully mapped (Cook et al., 2019). The two sexes have 294 common neurons, 8 hermaphrodite-specific neurons, and 91 male-specific neurons. Even the common neurons display sexual dimorphisms at molecular and circuit levels (Portman, 2017). Males and hermaphrodites elicit various behavioral differences, including pheromone preference (White et al., 2007; Fagan et al., 2018), food searching (Lipton et al., 2004; Ryan et al., 2014), and complex cue decision-making and learning (Sammut et al., 2015; Tanner et al., 2022). However, how these male-specific behaviors change with age is less well studied; previous neuronal aging studies have focused primarily on hermaphrodites (Kauffman et al., 2010; Arey et al., 2018). Male mating ability declines early in adulthood due to increased muscle excitability (Guo et al., 2012), but how male cognitive functions change with age is less well understood. While whole-animal transcriptomic studies of the male have been done (Kim et al., 2016; Ebbing et al., 2018; Reilly et al., 2021), less is known about male neuronal transcriptome changes with age.

In this study, we characterized male neuronal aging phenotypes. We found that male *C. elegans* can carry out positive Pavlovian olfactory learning and memory, and this ability declines with age. Similarly, sex-specific pheromone chemotaxis behavior, which is stronger in males than in females (White et al., 2007), also declines with age, as does sensory neuron morphology. To determine how gene expression changes may contribute to these age-related changes, we performed neuron-specific RNA-sequencing of Day 1 and Day 8 adult males. We found that metabolic and GPCR expression changes with age in males more than in similarly-aged hermaphrodites. We also found that epigenetic regulators are upregulated specifically in the aged hermaphrodite X chromosome. Lastly, we identified a gene that declines with age specifically in males, the aspartic protease *hrg-7*, and found that this gene is required for male cognitive functions. Our work suggest that transcriptomic changes in neurons with age contribute to the behavioral and morphological declines exhibited in aging males.

## Results

### Male neuronal aging in behavior and neuron morphology

Since wild-type males raised together exhibit shortened lifespans due to male pheromone (Shi et al., 2017), we used a pheromone-production defective mutant, *daf-22*, to measure normal aging in males, in a background that produces high incidence of males (*him-8*). We find that *him-8;daf-22* males have a lifespan similar to hermaphrodites from the same strain, which indicates the strain is appropriate for aging experiments (Figure 1A).

**Figure 1.**
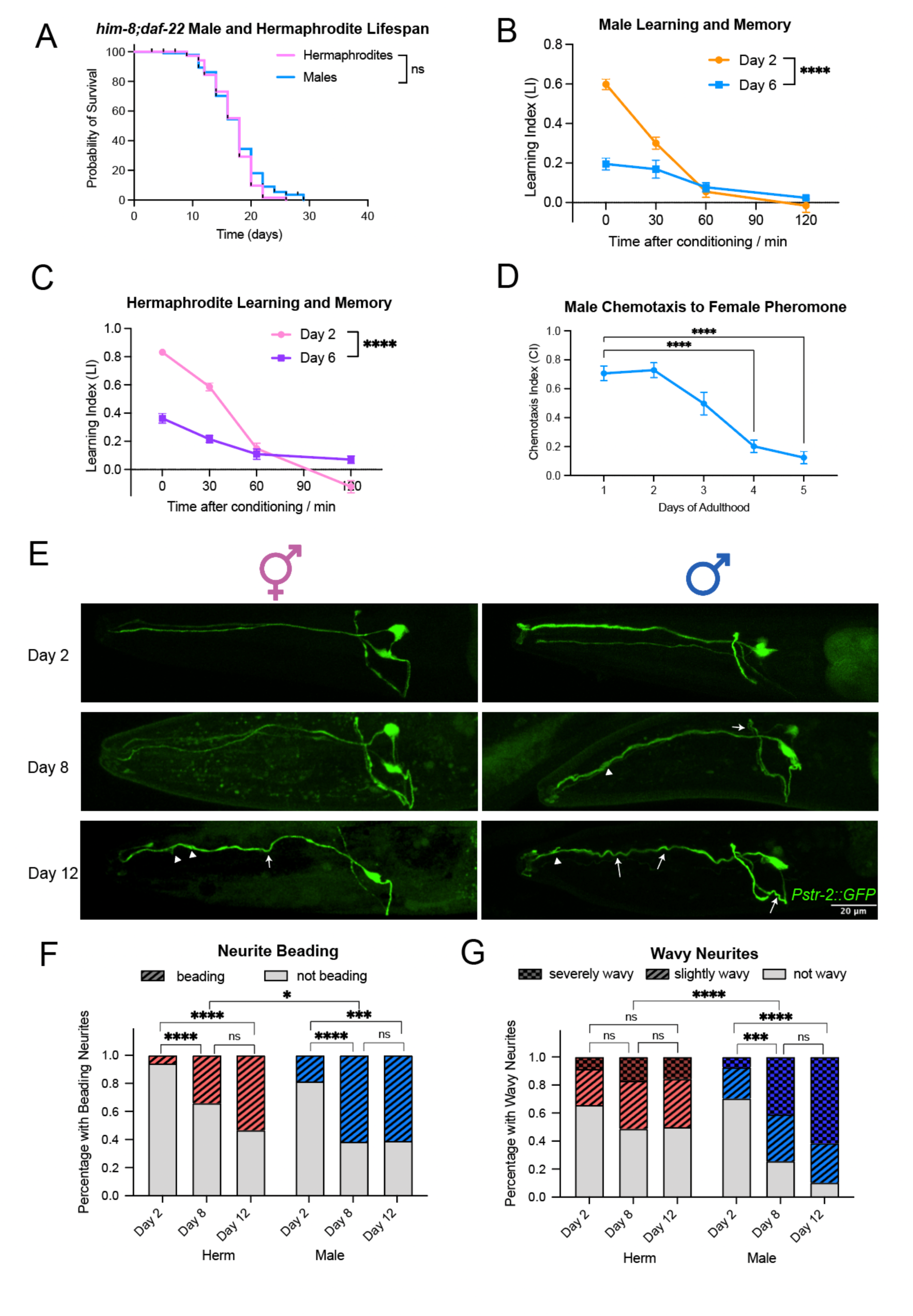
Male Neuronal aging can be characterized through aging phenotypes. (A) Lifespan of him-8;daf-22 males and hermaphrodites. ns: p > 0.05, Gehan-Breslow-Wilcoxon test. (B) Male learning and short-term associative memory (STAM) ability is high in young worms and decrease in aged worms. (C) Hermaphrodite learning and STAM ability is high in young worms and decrease in aged worms. Chemotaxis index at 0min after conditioning measures learning, and 30min, 60min, and 120min measures short-term memory trajectory. N = 4 biological replicates, 5 chemotaxis plates per biological replicate. ****: p <0.0001, two-way ANOVA with Tukey post-hoc analysis. (D) Male chemotaxis toward pheromone decreases with age. Chemotaxis index at different ages measures male preference for female pheromone. N = 3 biological replicates, 5 chemotaxis plates per biological replicate. ****: p <0.0001, *: p<0.05, ns: p>0.05, two-way ANOVA with Tukey post-hoc test. (E) Representative images of AWC neuron morphology at Day 2, Day 8 and Day 12. Herm: hermaphrodites. Arrow: wavy neurites. Arrowhead: Beading and branching. (D) Quantification of neurite beading and wavy neurite phenotypes in both sexes. N = 35,41,45,27,39,41 respectively. ****: p < 0.0001, **: p<0.01, *: p <0.05, ns: p>0.05. Chi-square test.

Next, we assessed male learning and memory using an assay of Pavlovian association between food and butanone (Torayama et al., 2007; Kauffman et al., 2010). On Day 2 of adulthood, males demonstrated intact appetitive learning and memory functions that lasts for about 60 minutes (Figure 1); however, there was a significant decline in these abilities by Day 6 (Figure 1B), which is comparable to that observed in hermaphrodites (Figure 1C). This suggests that young males have cognitive functions that are similar to those of hermaphrodites, and that male neuronal aging takes place on a similar trajectory.

*C. elegans* males display a preference for both hermaphrodite and female ascaroside pheromones (White et al., 2007; Srinivasan et al., 2008), but whether this ability declines with age had not been characterized. Using *C. remanei* (true female) pheromones, we investigated males’ ability to chemotaxis towards these pheromones as they age. Day 1 males have a strong preference towards the female pheromone. However, this attraction declines with age. By Day 4, males lose their ability to chemotaxis towards the female pheromone, coinciding with the timeframe of their reduced mating capability (Lipton et al., 2004; Chatterjee et al., 2013) (Figure 1D).

Next, we characterized morphological changes that accompany the observed behavioral decline with age. The neurons that have been previously analyzed for morphological changes with age are motor neurons (Herndon et al., 2002; Pan et al., 2011; Tank et al., 2011; Toth et al., 2012), but here we are interested in examining the changes that may accompany loss of learning and behavior, which are mediated by sensory neurons (Kauffman et al., 2010; Lakhina et al., 2015; Arey et al., 2018). AWC neurons are chemosensory neurons responsible for butanone sensation, and we found that these neurons are required for short-term associative memory (Stein and Murphy, 2014; Arey et al., 2018). In both hermaphrodites and males, AWC neurons exhibit an increase in neurite beading and wavy neurites as they age. Notably, the severity of neurite beading and the wavy neurites in aged males was greater than that observed in age-matched hermaphrodites (Figure 1E-G). These findings suggest that neuronal aging in males presents some unique, sex-specific features when compared to hermaphrodite aging.

### The male neuronal aging transcriptome

To understand the molecular mechanisms that may underlie the behavioral changes with age that we observed, we performed neuron-specific sequencing on Day 2 and Day 8 male neurons. Through filtering separation, we are able to separate males and hermaphrodites from a mixed sample, and achieved >95% purity of male worms in our samples, thus making it possible for assessment of the male transcriptome. PCA of our neuronal samples showed a clear separation of the young and aged male neurons (Fig. 2A), suggesting that age-related differences were the dominant source of differential gene expression, and the batch effects of the samples are small. Differential expression analysis using DESeq2 identified 763 genes that were more highly expressed in young than aged male neurons and 1,063 genes that were more highly expressed in aged than young male neurons (log2(fold-change) >1.0 or <-1.0 and p-adj <0.05) (Figure 2B). A tissue-specific query showed that these differentially expressed genes had high prediction scores for the nervous system and neurons, validating that our differential expression dataset is neuron-specific (Figure S1).

**Figure 2.**
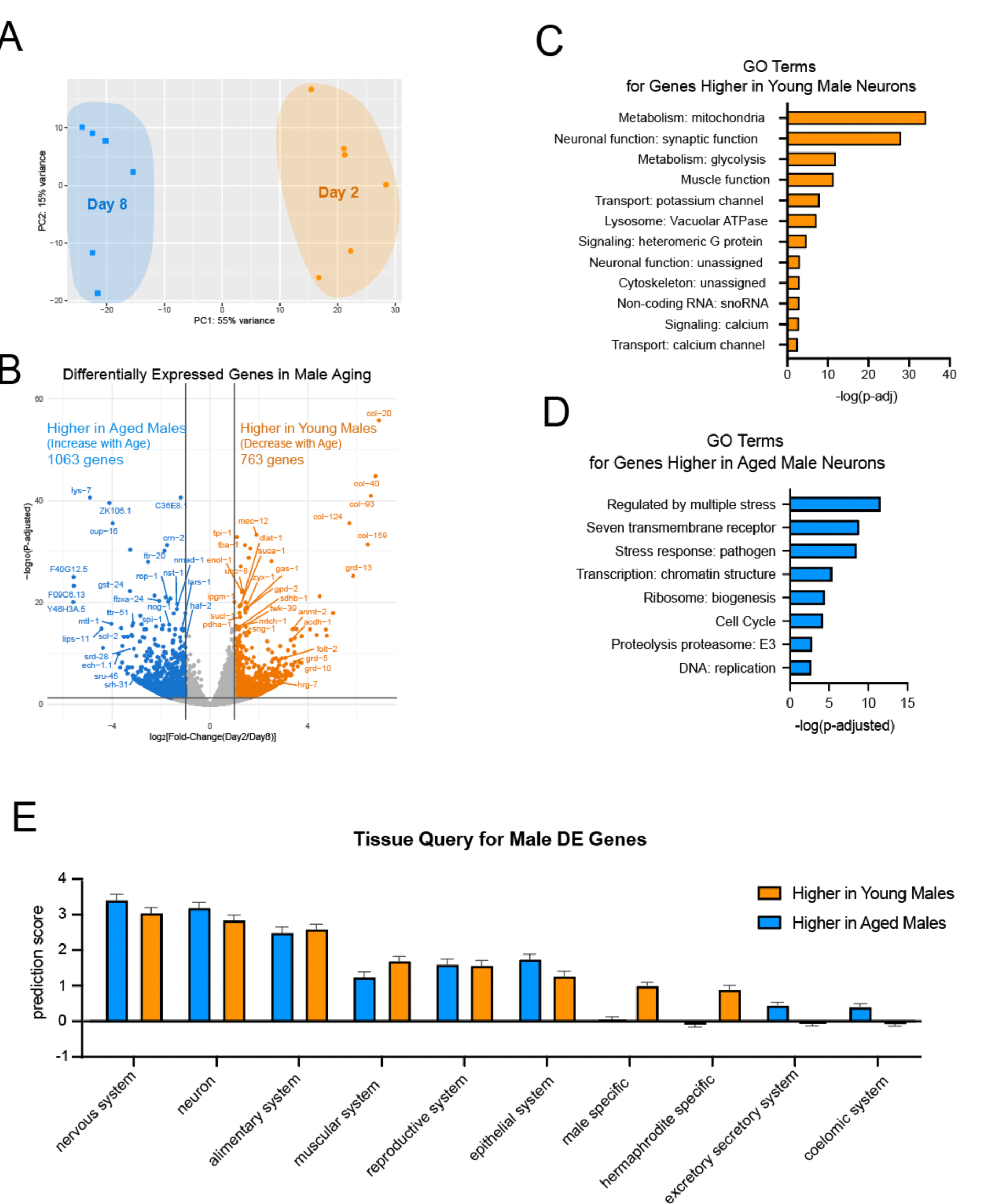
Male neuron sequencing identified gene expression change with age. (A) PCA plot shows age-related separation between young and aged male neuron collections. PCA plot generated using the DESeq2 package. (B) Volcano plot showing differentially expressed genes in male aging. Log2(fold-change) >1.0 or <-1.0, p-adj <0.05 are considered differentially expressed. Top differentially expressed genes are labeled. (C) Gene Ontology results from genes significantly more highly expressed in young male neurons compared with aged male neurons. Gene ontology results from Wormcat 2.0 category 2. (D) Gene Ontology results from genes significantly more highly expressed in aged male neurons compared with young male neurons. Gene ontology results from Wormcat 2.0 category 2. (E) Tissue query for male differentially expressed genes. Top 500 genes more highly expressed in young male neurons or aged male neurons are input for the tissue query. Only prediction scores for major tissues and neuron is shown. Prediction scores and SEM is generated from the tissue query website.

Next, we used Gene Ontology (GO) analysis to assess predicted functions of differentially expressed genes with age. Genes more highly expressed in young male neurons were enriched in GO terms related to mitochondrial metabolism (e.g., *idhg-2, gpd-2, ctc-2*), neuronal synaptic function (e.g., *nlp-20, rab-3, dat-1*), signaling and transport (e.g., *egl-20, twk-30, ccb-2*), and structural and cytoskeletal proteins (e.g., *col-20, col-40, tnt-2*) (Figure 2C). The age-related decline in expression of these genes that are essential to neuronal cell function correlates with the behavioral and morphological declines we noted in aging males.

By contrast, genes higher in aged neurons are enriched in GO terms for stress response (e.g., *mtl-1, cyp-14A5, lys-7*), transmembrane serpentine receptors (e.g., *sru-45, srh-31, srd-28*), and transcription regulation (e.g., *eef-2, his-42, set-32*) (Figure 2D). The increased expression of these genes may suggest transcriptional dysregulation and a response to increased stress in aged neurons.

### Comparison with hermaphrodite neuron-specific sequencing reveals sex-specific features

To understand the sex-specific differences between hermaphrodite and male neuronal aging, we compared young and aging hermaphrodite neurons (Weng et al., 2023) to our newly-generated young and aged male neuronal transcriptomes. Samples from males and hermaphrodites distinctly separate along PC1, while PC2 captures age-induced variations (Fig. 3A); biological replicates clustered together, indicating the differences we see in expression were not due to sample collection artifacts. We note that the hermaphrodite young and aged samples showed a greater separation than the male samples. This difference may be due to the difference in collection age for the neurons: while hermaphrodite young neurons were sorted on Day 1, male neurons were sorted on Day 2 due to the filtration protocol. To control for the age-related differences, we included a Day 2 hermaphrodite neuron collection, and this sample demonstrated a PC2 value comparable to that of the male Day 2 neuron samples (Figure 3A).

**Figure 3.**
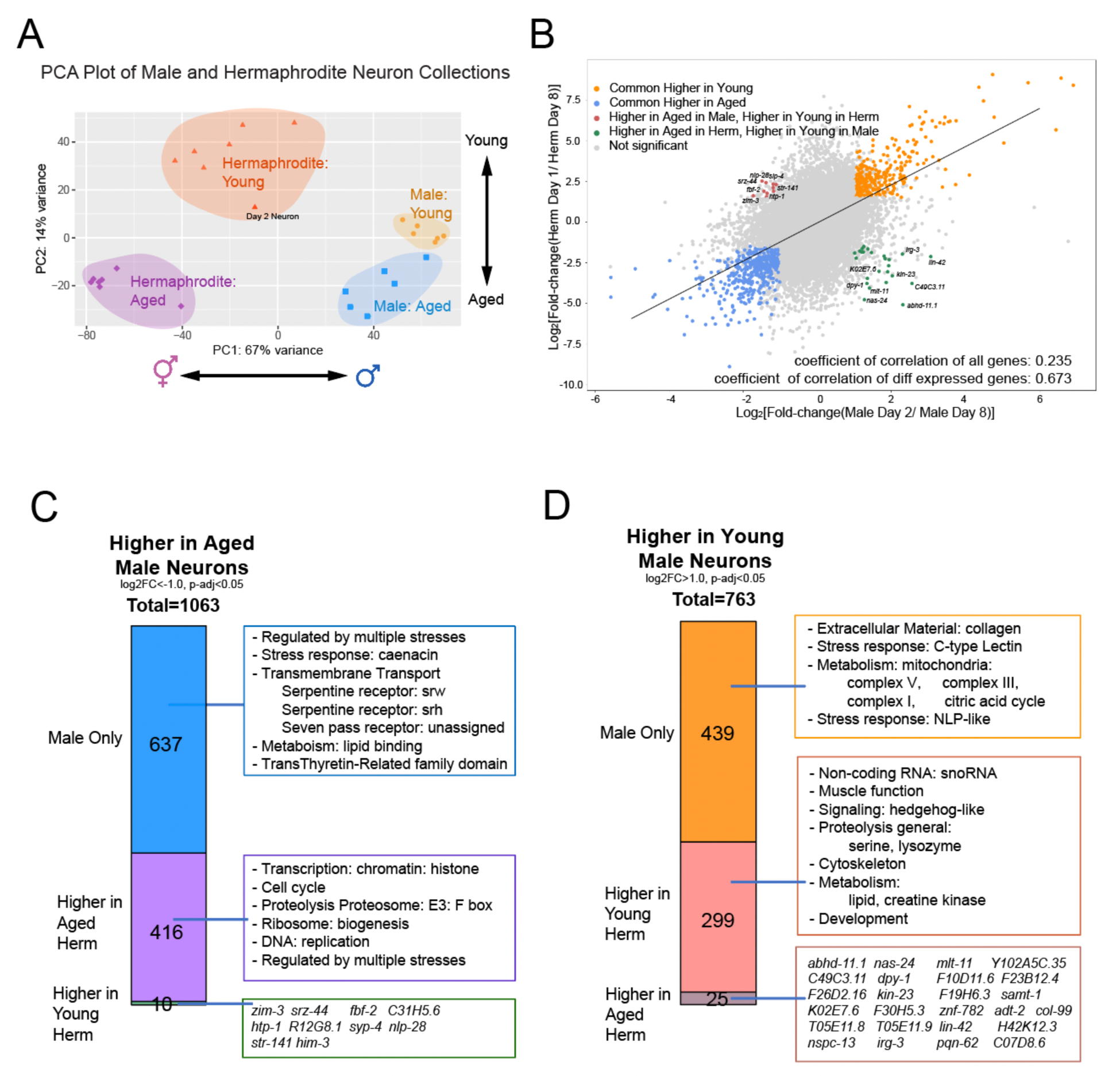
Sex-specific dimorphism in the neuronal aging transcriptome. (A) PCA plot of hermaphrodite and male neuron collection samples. 6 replicates of wild-type hermaphrodites and 1 replicate of *him-8;daf-22* hermaphrodite (Labeled as Day 2 neuron) is shown. The plot is generated from the DESeq2 package. (B) Correlation of log2[fold-change(young/aged)] in hermaphrodites (y-axis) and males (x-axis). Differentially expressed gene cutoff is log2[fold-change(young/aged)] > 0.5 or < -0.5, p-adj <0.05. Genes differentially expressed are color-labeled as shown. (C-D) Fraction of male genes more highly expressed in young neurons (C) and aged neurons (D) that are either only differentially expressed in males, or higher expressed in young neurons in hermaphrodites, or higher expressed in aged neurons in hermaphrodites. Herm: hermaphrodites. Gene ontology results from WormCat 2.0 category 2.

Next, we analyzed the correlation of young vs aged differential gene expression fold-change in hermaphrodites and males (Figure 3B). The correlation between the significantly differentially-expressed genes in hermaphrodites and males was high, while the genes not significantly differentially-expressed had a lower correlation. This suggests that hermaphrodites and males share many age-related gene expression changes, forming a conserved aging process across both sexes (Figure 3B, orange and blue). Although highly correlated, there is a small set of genes that are differentially expressed in the opposite direction between hermaphrodites and males (red, green on plot). Some genes show higher expression in young hermaphrodite neurons but are increased in aged male neurons, such as *zim-3, srz-44,* and *fbf-2*. Conversely, *lin-42, irg-3,* and *kin-23* among others exhibit higher expression in young male neurons and aged hermaphrodite neurons (Table 1).

Although some enriched GO terms are shared between aging male and hermaphrodite neurons (including transcription, cell cycle, proteostasis, stress responses, ribosome biogenesis, and DNA replication), there were a few notable notable differences (Figure 3C-D). Genes with a higher expression in young neurons in both sexes are enriched in cytoskeletal organization, proteolysis, and lipid metabolic functions. However, genes that are elevated in young male neurons but not in hermaphrodite neurons are associated with mitochondrial metabolism, including Complex I, III, IV, and citric acid cycle genes (Figure 3C). This suggests a decline in cellular respiratory and energy-generating processes, particularly in male neurons as they age. Conversely, genes that exhibit higher expression with age in both sexes show enrichment in proteolysis proteasome and epigenetic modification pathways, but genes that are only upregulated in aged male neurons additionally demonstrate a marked enrichment in seven-transmembrane receptors, most of which are serpentine receptors (Figure 3D).

### Neuronal functional genes are higher in young neurons in both sexes

Very few neuronal genes are upregulated with age (Figure 4A, S2A), but among those were genes encoding two neuropeptides (*nlp-34* and *nlp-29*) and OSM-10, which is known to be expressed in sensory neurons in the hermaphrodite.

**Figure 4.**
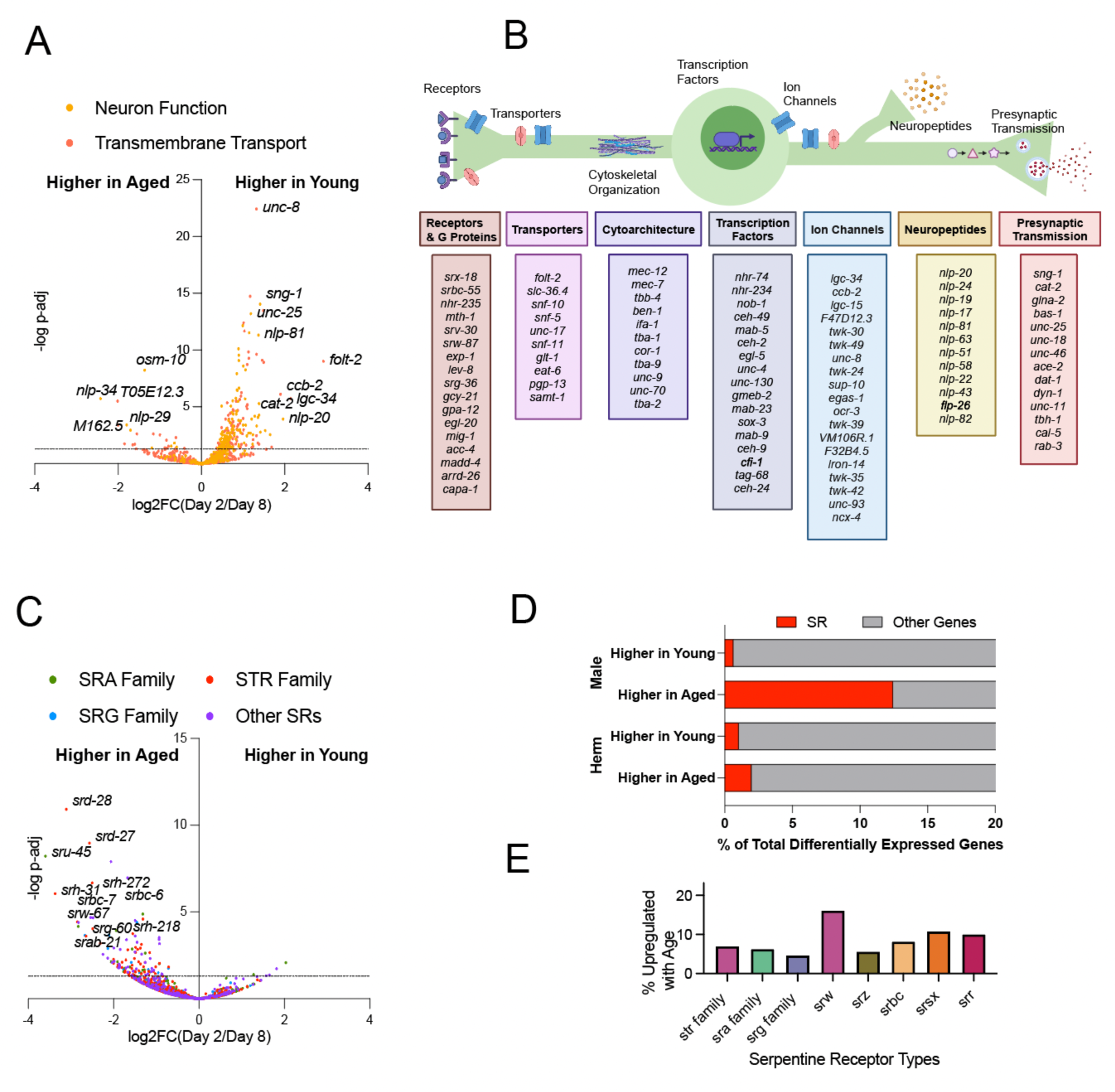
Neuronal genes are differentially expressed with age in males. (A) Volcano plot of neuron functional genes and transmembrane transport genes’ expression change with age in males. (B) Neuron function and transmembrane transport genes that downregulate with age in males play various roles in a neuron. (C) Volcano plot of serpentine receptor (SR) genes’ expression change with age in males. (D) Percentage of SRs that are differentially expressed with age in males and hermaphrodites. (E) Percentage of SRs in each SR family that are upregulated with age in males.

Perhaps unsurprisingly, we found that neuronal functional genes for the most part decline with age in both hermaphrodite and male neurons. These neuronal genes participate in various aspects of neuronal function, including signaling, transport, transcription, and cytoskeletal architecture (Figure 4B). Upregulated in young genes include postsynaptic receptors and G protein signaling pathway genes, such as the *srx-18, srbc-55, srv-30, srw-87, srg-36, Methuselah/mth-1,* and *capa-1* GPCRs (although the majority of serpentine receptors are upregulated with age - see below and Figure 4C), nuclear hormone receptor *nhr-235*, neurotransmitter receptors *exp-1, lev-8,* and *acc-4*, receptor tyrosine kinase *egl-20*, WNT receptor *mig-1*, axon guidance receptor *madd-4*, and the heteromeric G proteins *gpa-12, gcy-21*, and *arrd-26*. Transmembrane porters and ion channels are also more highly expressed in young male neurons, including potassium channels *twk-30, twk-49, F47D12.3, twk-24, twk-39, sup-10, VM106R.1, F32B4.5, unc-93, twk-35, twk-42,* and *lron-14*, calcium channels *ccb-2, ocr-3, ncx-4*, sodium channels *unc-8, egas-1*, ligand gated channels *lgc-34, lgc-15*, amino acid transporters *slc-36.4, snf-10*, *glt-1*, GABA transporter *snf-5, snf-11*, ATPase coupled transporter *pgp-13*, molybdate ion transporter *samt-1*, and sodium:potassium-exchanging transporter *eat-6*.

There are also genes that regulate the cytoskeletal architecture that decline with age, including tubulins *mec-12, mec-7, tbb-4, ben-1, tba-1, tba-9,* and *tba-2*, intermediate filament *ifa-1*, and actin filament binding proteins *cor-1, unc-9,* and *unc-70.* Neuronal transcription factors *ceh-2, egl-5, unc-4, unc-130,* and *ceh-4*, and neuropeptides, including *nlp-20, nlp-24, nlp-19,* and *nlp-17* also decline with age.

Synaptic proteins, including synaptic vesicle formation and release proteins *sng-1, unc-18, unc-46, dyn-1, unc-11, cal-5, rab-3*, and neurotransmitter synthesis and uptake enzymes *cat-2, glna-2, bas-1, unc-25, ace-2, dat-1*, and *tbh-1* also decline with age.

Overall, this wide range of neuronal genes being downregulated with age may predispose the neurons to be unable to fulfill various cellular processes such as gene regulation, structural integrity, signal propagation, and synaptic transmission, thus leading to the behavioral and morphological changes we see in aged animals.

### Chemosensory GPCRs are upregulated in aged neurons in males

We also found that chemosensory GPCRs, especially serpentine receptors, were upregulated with age in male neurons (Figure 4C). In fact, about 10% of genes that were upregulated with age were serpentine receptors (Figure 4D). This is a phenomenon specific to males: serpentine receptors are not similarly enriched in genes that upregulate in aged hermaphrodite neurons (Figure S2C). The serpentine receptors that are upregulated with age include all major serpentine receptor classes, including *str*, *sra*, and *srg* super family genes, *srw, srz, srbc, srsx*, and *srr* families. The differential expression of these serpentine receptors, which plays a crucial role as chemosensory GPCRs, may underlie the behavioral differences of odorant sensation and chemotaxis in males and hermaphrodites (Portman, 2007; Loxterkamp et al., 2021).

### Canonical metabolic genes are downregulated with age in male neurons

One category of genes that decline specifically in aging male neurons but not in aging hermaphrodite neurons are those involved in canonical energy-generating respiratory metabolic pathways. Mitochondrial-related genes are more highly expressed in young male neurons (Figure 3D, 5A): glycolytic and mitochondrial metabolic genes are enriched in genes more highly expressed in young male neurons. We found that several enzymes of that mediate glycolysis are downregulated with age at least 2-fold (labeled in red), including 6-PhosphoFructoKinase *pfk-1.1*, 6-PhosphoFructoKinase *aldo-1*, glyceraldehyde-3-phosphate dehydrogenase *gpd-2*, phosphoglycerate mutase *ipgm-1*, phosphoglycerate mutase *enol-1*, and pyruvate dehydrogenases *pdha-1, pdhb-1, dlat-1, dlat-2,* and *dld-1*. Mitochondrial metabolic genes are also significantly downregulated, including TCA cycle enzymes *idhg-2, dlst-1, dld-1, sucl-1, suca-1, sdha-1, sdhb-1, sdhd-1,* and *mdh-1*, and electron transport chain complex subunits for each complex. This includes Complex 1 (ubiquinone) genes *gas-1, nduo-3, Y54F10AM.5, Y63D3A.7, F31D4.9, lpd-5, nduo-6, C33A12.1, Y71H2AM.4, C18E9.4,* and *nduo-4*; Complex II (succinate dehydrogenase) genes *sdha-1, sdhb-1,* and *sdhd-1*; Complex III (Coenzyme Q – cytochrome c) and cytochrome C genes *F45H10.2, cyc-1, ucr-11, ctb-1, ucr-2.2, T02H6.11,* and *C14B9.10;* Complex IV genes, *ctc-2, cox-6C, cox-4,* and *cox-6b*; and ATP synthesis subunits *asb-2, R04F11.2, asg-2, Y82E9BR.3, Y69A2AR.18, atp-2, atp-1,* and *R53.4*. The downregulation with age of these fundamental carbohydrate metabolic genes may impose a sex-unique challenge to male neuronal metabolism in aged animals.

**Figure 5.**
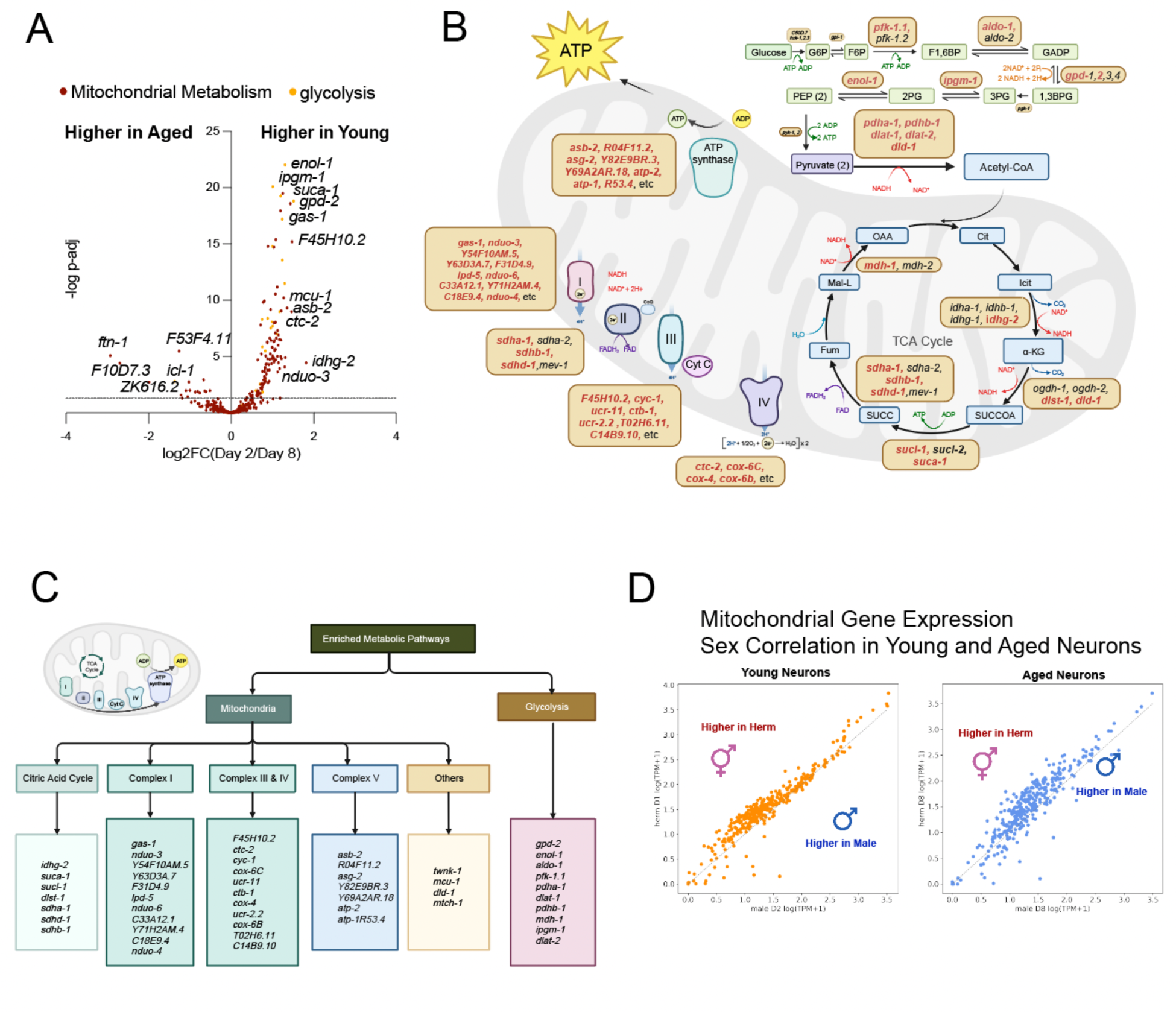
Mitochondrial genes are differentially expressed with age in males. (A) Volcano plot of mitochondrial metabolic and glycolysis genes’ expression change with age in males. (B) Metabolic flux of glycolysis, TCA cycle and electron transport chain. Metabolic enzymes whose gene expression is downregulated with age in male neurons are labeled in red. (C) Metabolic genes that are more highly expressed in young neurons participate in different metabolic categories. (D) Correlation of transcript per million (TPM) number mitochondrial metabolic genes in hermaphrodite neuron and male neuron in young and aged animals. Dots above the dashed line indicates expression level is higher in hermaphrodite neurons than male neurons.

To understand whether this sex-specific downregulation of metabolic genes is due to young males having higher expression of metabolic genes than hermaphrodites, or due to aged males expressing lower levels of metabolic genes, we compared the correlation of transcript number per million transcripts (TPM) of metabolic genes between hermaphrodites and males (Figure 5D). We found that in young animals, males and hermaphrodites have a good correlation of mitochondrial-related gene expression (although hermaphrodites have slightly higher expression), but hermaphrodites have even higher metabolic gene expression than males in aged animals. Thus, the decline of metabolic genes in aged animals is not due to a high expression of metabolic genes in young animals, but a decline in expression of metabolic genes in aged animals, thus very likely compromising their energy production and cellular activity maintenance.

### X chromosomal genes exhibit changes with age

The sex chromosome (X) contributes genetically to sex differences; thus, we focused on the X chromosomal changes during neuronal aging of hermaphrodites and males. In *C. elegans*, hermaphrodites possess two X chromosomes (XX), whereas males have just one (XO) (Madl and Herman, 1979). To counteract this imbalanced chromosome number, worms carry out dosage compensation mechanisms to ensure a balanced expression of X chromosomal genes across both sexes (Meyer and Casson, 1986). Dosage compensation should be maintained throughout life to achieve normal gene expression levels. Therefore, we analyzed the expression of all of the chromosomes with age, and further, expression of X chromosome genes in both males and hermaphrodites. We found that in all autosomes, there is male-biased gene expression in young neurons, meaning genes on that chromosome are more likely to be higher expressed in males than hermaphrodites, and this male bias declines in aged neurons (Figure 6A, S3).

**Figure 6.**
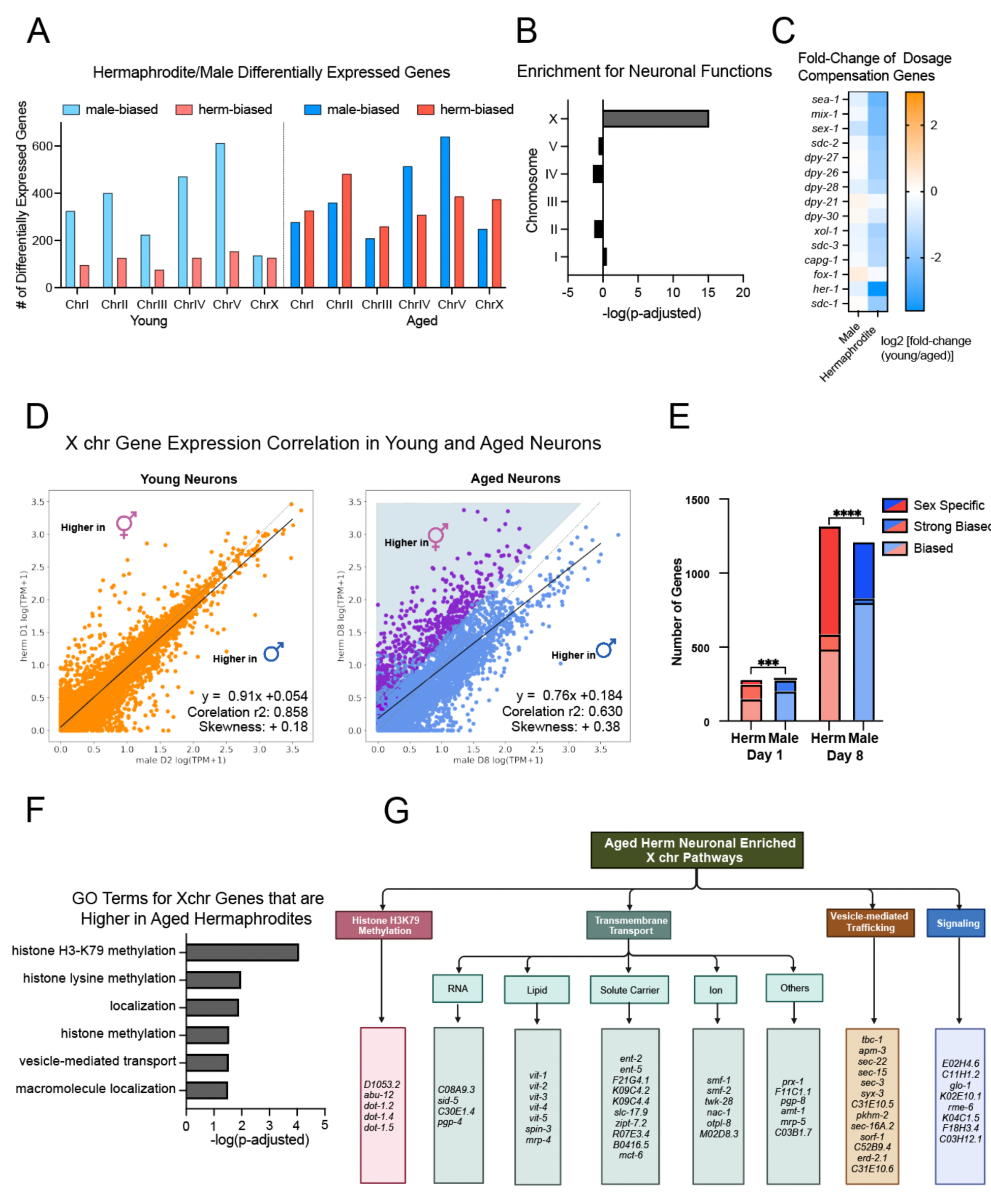
X chromosomal gene regulation exhibits sex-specific dimorphisms. (A) The number of genes significantly expressed between sexes on each chromosome. Male-biased genes are significantly more highly expressed in male neurons compared with aged-controlled hermaphrodites, and hermaphrodite-biased genes vice versa. Genes significantly expressed are have a log2[fold-change(herm/male)] > 2 or < -2, p-adj < 0.05. (B) Genes from the X chromosome are enriched in neuronal functions. Gene ontology of all protein-coding genes in a chromosome is used as input for WormCat gene ontology analysis. (C) log2[fold-change(young/aged)] of dosage compensation complex genes. (D) Male-hermaphrodite correlation of gene expression on the X chromosome. In young neurons, gene expression correlation between 2 sexes is higher than in aged neurons. Linear regression fitted function, Pearson’s correlation r2, and distribution skewness are calculated using Python. Genes significantly higher expressed in aged hermaphrodites compared with aged males are labeled in purple. (E) Number of genes that are sex-specific (TPM > 1 in one sex, and TPM < 0.01 in the other sex), sex-biased (TPM > 0.01 in both sexes, and TPM ratio > 2, and p-adj < 0.05) and strongly sex-biased (TPM > 0.01 in both sexes, and TPM ratio > 8, and p-adj < 0.05). ***: p < 0.001, ****: p<0.0001, chi-square tests. (F) Gene ontology terms for X chromosomal genes that are higher expressed in aged hermaphrodites. GO Terms measured using gProfiler. (G) Genes belonging to the enriched GO terms from (F).

Among all the chromosomes, the X chromosome has the highest enrichment of neuron function genes, indicating it as a potentially vital regulator in cognitive aging, regardless of sex (Figure 6B). However, we observed an age-related upregulation of dosage compensation complex genes only in hermaphrodite neurons (Figure 6C). This suggests a potential dysregulation of the dosage compensation/X inactivation process in aged hermaphrodites, which may contribute to sex-specific changes with age in X chromosomal genes only in hermaphrodites.

To test that hypothesis, we plotted the correlation of X chromosomal gene expression in both young and aged neurons. Previously, Albritton et al. found that genes on the X chromosome are more likely to be hermaphrodite-biased in their expression, although in general sex-biased genes are more likely male-biased (Albritton et al., 2014). In our study, we found that in young neurons, the gene expression levels between hermaphrodite and male neurons correlate well (r^2^ = 0.85), suggesting a stable gene expression with minimal sex-specific variances (Figure 6D). However, in aged neurons, this correlation decreased greatly (r^2^ = 0.63), indicating a sex-specific age-dependent differential regulation of X-chromosomal genes. Notably, the expression distribution is slightly skewed (skewness = +0.38), with more genes having higher expression in aged hermaphrodites than aged males (Figure 6D, Figure S2D).

We next quantified the number of X chromosome genes that are slightly (2-fold difference), strongly (8-fold difference) or completely sex-specific (expressed only in one sex). We found that on the X chromosome, there are more sex-biased genes expressed in aged neurons than in young neurons, and aged hermaphrodites more strongly express neuronal X chromosome sex-biased and sex-specific genes than do males (Figure 6E).

GO term analysis of the X-chromosomal genes that are significantly more highly expressed in aged hermaphrodites revealed an enrichment in pathways like histone H3K79 methylation and localization (Figure 4F). The “localization” gene ontology term included genes that function in (1) transmembrane transport, including mediating the transportation of RNA (e.g., *sid-5, pgp-4*), lipid (*e.g., vit-1, spin3, mrp-4*), ions (e.g., *smf-1, smf-2, twk-28*), and solute carriers (e.g., *ent-2, ent-5, slc-17.9*), and others (e.g., *prx-1 pgp-8, amt-1*), (2) vesicle-mediated transport, including in exocytosis (e.g., *sec-15, sec-3, syx-3*), endocytosis (e.g., *apm-3, sorf-1*), and ER/Golgi trafficking (e.g., *sec-22, erd-2.1, pkhm-2, sec16A.2*) (3) signaling, including receptors (e.g., *E02H4.6, C11H1.2*), and G proteins (e.g., *glo-1, rme-6, K02E10.1*) (Figure 4G).

The changes in these X chromosomal genes may contribute to the sexual dimorphic phenotypes we see in aged animals. Histone H3K79 methylation genes may contribute to chromatin regulation of enhancer regions and gene expression(Esse et al., 2019; Esse and Grishok, 2020), while neuronal transport and localization genes may contribute to ion transport and vesicular signaling. Therefore, these X chromosomal differentially-expressed genes may play roles in the observed sexual dimorphism in aging behaviors and transcriptomes.

### The Heme Responsive Gene *hrg-7* may be required for young male behaviors

We would hypothesize that genes differentially expressed with age may play an important role in cognitive decline. We tested several genes that significantly decline with age in males but do not decline with age in hermaphrodites (Figure S4). *hrg-7,* which encodes an aspartic-type endopeptidase, is more highly expressed in young male neurons than in aged male neurons (Figure 2B), and its expression declines more significantly with age in males compared to hermaphrodites (Figure 7A). This was somewhat surprising, as HRG-7 is expressed in the intestine as well as the amphid and phasmid neurons, and is secreted as an inter-organ signaling protein in hermaphrodites, sending signals from the intestine to the neuron; HRG-7 is thought to be crucial for the regulation of heme homeostasis (Sinclair et al., 2017).

**Figure 7.**
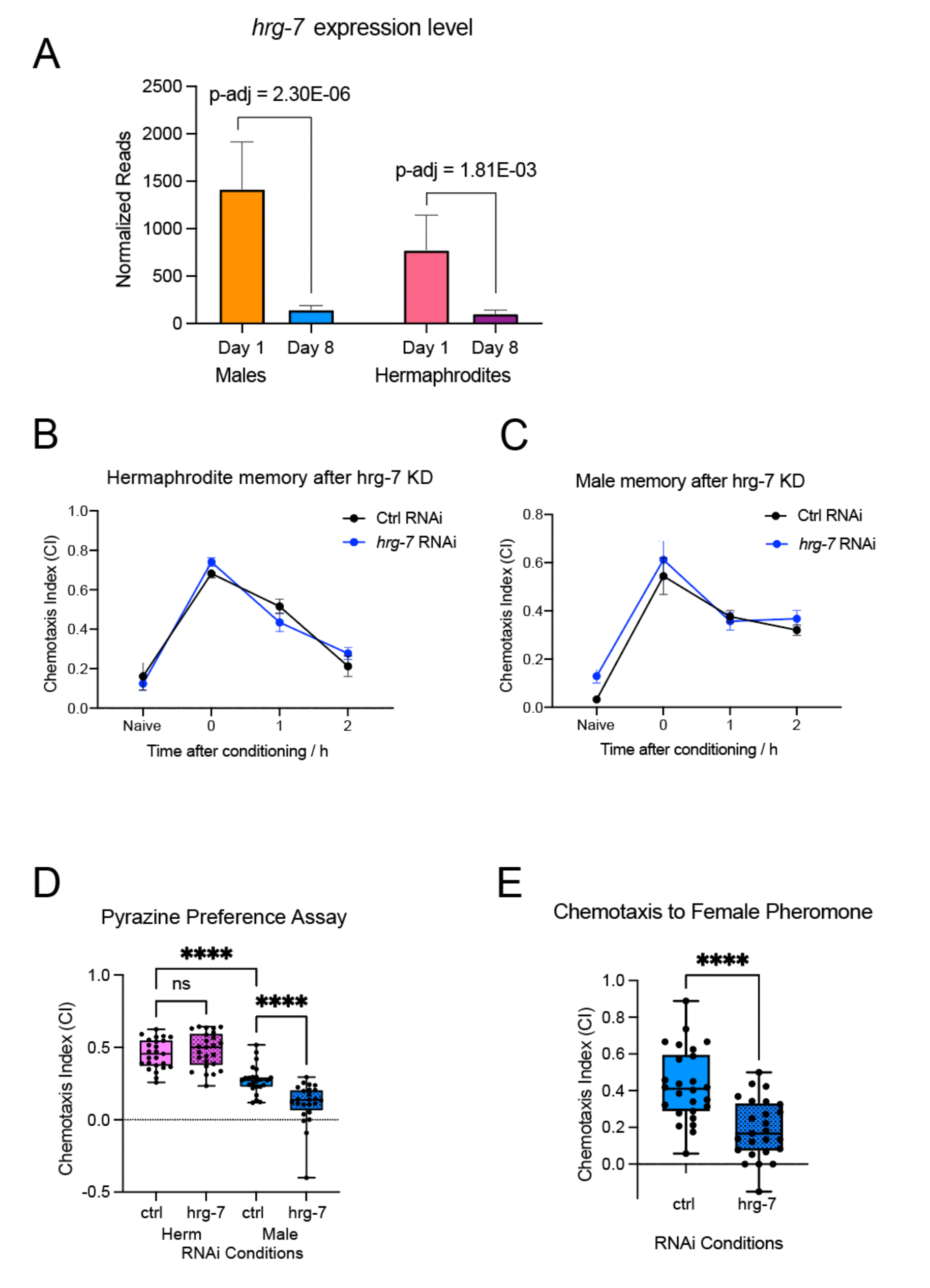
*hrg-7* is required for male-specific chemotaxis functions. (A) *hrg-7* expression level in males and hermaphrodites. *hrg-7* is significantly downregulated with age. P-adjusted value from DESeq2. (B) Hermaphrodite learning and STAM ability under control and *hrg-7* RNAi. (C) Expression of *hrg-7* in Day 2 hermaphrodites and males of IQ7701 strain. *hrg-7* expression is identified in male tail neurons. (C) Male learning and short-term associative memory (STAM) ability under control and *hrg-7* RNAi. Chemotaxis index at 0min after conditioning measures learning, and 30min, 60min, and 120min measures short-term memory trajectory. N = 4 biological replicates, 5 chemotaxis plates per biological replicate. ****: p <0.0001, two-way ANOVA with Tukey post-hoc analysis. (E) Pyrazine chemotaxis is compromised in *hrg-7* neuron-specific RNAi knockdown worms, but only in males. N = 4 biological replicates, 25 chemotaxis plates total. ****: p < 0.0001, ns: p > 0.05. One-way ANOVA with Dunnet post-hoc test. (F) Male’s ability to chemotaxis to female pheromone is compromised in *hrg-7* neuron-specific RNAi knockdowns. RNAi is performed in the TU3595 neuron-specific RNAi strain. N = 5 biological replicates, 5 chemotaxis plates per replicate.

To test whether HRG-7 has any roles in male-specific behavior, we knocked it down in both sexes and tested memory and chemotaxis. While the reduction of *hrg-7* does not affect learning and memory in either sex (Figure 7B, C), we found that *hrg-7* RNAi reduced pyrazine chemotaxis specifically in males (Figure 7D). Furthermore, *hrg-7* knockdown in male neurons impairs their ability to chemotaxis to female pheromone (Figure 7E). These results indicate a role for HRG-7 specifically in regulating male behaviors, and suggest that its age-related decline may contribute to the reduced chemotaxis and pheromone sensing behavior we observed in aged males. This discovery further strengthens the connection between sex-specific transcriptomic changes and behavioral decline during aging. Our findings indicate that we can use this dataset to identify genes that contribute to sex-specific phenotypes during cognitive aging, and it will offer valuable insights into sex-specific individualized interventions to regulate cognitive decline in both sexes.

## Discussion

Here, we characterized behavioral and morphological changes during male cognitive decline, discovering that male learning and memory function, pheromone preference, and male neuronal morphology integrity all decline with age. We then performed neuron-specific sequencing on young and aged male neurons to identify gene expression changes. We found that neuronal and mitochondrial metabolic genes decrease with age, and some GPCRs and stress-response-related genes increase with age. We then performed a comparison with the hermaphrodite neuron transcriptome both at the genome-wide and X chromosomal levels to identify sexual dimorphic gene expression features with age. Through our sequencing and analysis, we discovered genes that regulate behavioral change with age in a sex-specific manner, linking transcriptomic analysis with behavioral outcomes.

We found that some male neuronal aging phenotypes are more severe than is observed in hermaphrodites. While male learning and short-term memory decline are comparable to that of hermaphrodites, the male’s ability to chemotaxis to female pheromone declines starting Day 3, before many behavioral deficits with age observed in hermaphrodites. Males also have a more severe wavy neurite phenotype compared with hermaphrodites in aged neurons. Notably, these phenotypic declines coincide with the timescale of male mating ability decline (Chatterjee et al., 2013). We hypothesize that the male cognitive decline can be explained in the evolutionary context. By adult Day 3, male mating potency is significantly decreased (Chatterjee et al., 2013), and there may be no survival advantage for individuals with improved cognitive ability post-reproduction. By contrast, hermaphrodites’ reproductive ability is maintained until Day 8-10 (Hughes et al., 2007; Luo et al., 2010), so there may be an evolutionary benefit to maintaining hermaphrodite-specific and sex-shared behaviors slightly longer with age.

We also identified male-specific transcriptomic changes with age. We found that metabolic genes, such as *acdh-1, abhd-11.1, idhg-2,* and *dhs-2*, decline with age in male neurons (Figure 2C, Figure 5). Nervous system metabolism is crucial to the execution of its functions. In humans, the brain consumes 20% of the total energy intake despite being only 2% of the total weight (Raichle and Gusnard, 2002), and females have increased mitochondrial activity and consume more oxygen than their male counterparts at the same age (Demarest and McCarthy, 2015). The human female brain also maintains metabolic youth better with age compared with the male brain (Goyal et al., 2019), which is consistent with our observations here in *C. elegans* neurons. Mitochondrial dysfunction is associated with various neurodegenerative diseases (Johri and Beal, 2012; Mor et al., 2020). Thus, this decrease in metabolic gene expression may pose a sex-specific threat to aged neurons in males.

In addition to genes with decreased expression with age, we also found that male neurons have increased GPCR expression with age, which is not observed in hermaphrodites. Interestingly, many of these GPCRs are not detected in hermaphrodites (e.g., *sru-45, srh-31, srg-60, srh-272*) (Taylor et al., 2021). The sex dimorphic expression of chemosensory GPCRs is not surprising given the chemosensory behavioral differences observed by us and others (Figure 1, Figure 6). The upregulation of these GPCRs with age, though, is more interesting. Cognitive aging can also be associated with neuronal hyperactivity (Bakker et al., 2012) and the expression of GPCRs could contribute to cellular excitability. Thus, the upregulation of GPCRs may promote neuronal hyperactivity and promote cognitive aging.

In addition to looking at the transcriptomic changes in general, we also focused on the sex chromosome. Here, we found that the X chromosome is enriched in genes with neuronal functions. These include neurotransmitter release and receptors (e.g. *acr-8, acr-12, dop-1, mgl-1*), signal transduction (e.g., *aex-3, mec-2, odr-10, osm-11*), and neuropeptides (e.g., *flp-2, flp-3, flp-5, nlp-1, nlp-2*). Previously, the X chromosome had been noted to express genes in a sex-biased manner (Albritton et al., 2014); here we further characterized how the X chromosomal gene expression bias in the nervous system changes with age. We found that in all autosomes, there is male-biased gene expression in young neurons, meaning genes on that chromosome are more likely to be higher expressed in males than hermaphrodites, and this male bias declines in aged neurons (Figure S3). However, the X chromosome is different from the autosomes, with a hermaphrodite-biased gene expression profile, and this bias is greater in aged neurons. We then examined individual genes with a hermaphroditic bias on the X chromosome, and found that the genes more highly expressed in hermaphrodites in aged neurons are enriched in H3K79 histone modification, in addition to neuronal signaling genes. In mammals the H3K79 methyltransferase DOTL1 mediates neurogenesis and neuronal energy metabolism (Appiah et al., 2023; Van Heesbeen et al., 2023). DOT1L in C. elegans has been shown to regulate H3K9me2 in enhancer regions and regulate RNAi efficiency through suppression of heterochromatin, thus eliciting a global impact on gene expression (Esse et al., 2019; Esse and Grishok, 2020). Interestingly, in mammals, histone modification enzymes are also among the X chromosomal inactivation (XCI) escapee genes that are found to be expressed from the inactivated X chromosome (Berletch et al., 2010). The expression of the XCI escapee gene, Kdm6a, in males has beneficial effects on cognitive aging (Shaw et al., 2023). Thus, X chromosomal hermaphroditic bias in gene expression may be a conserved feature, and these biased genes may act to regulate cognitive health with age in hermaphrodites and females. Together, our analyses uncovered genes and pathways that may affect neuronal aging in a sex-specific manner.

## Acknowledgments

We thank the *Caenorhabditis* Genetics Center (CGC) and the Hamza Lab for strains, WormBase (version WS289) for information, Jasmine Ashraf, William Keyes, and Titas Sengupta for help with experiments, members of the Murphy Lab for input on the manuscript, Christina DeCoste, Katherine Rittenbach and the Flow Cytometry Facility for cell sorting assistance, Wei Wang and the Genomics Facility for sequencing assistance, Lance Parsons for insights on data analysis, and Biorender.com for schematic design.

## Methods

### Worm maintenance

Strains used in this study:

CB1489: *him-8(e1489) IV*,

CQ675: *daf-22(m130) II; him-8(e1489) IV*,

CQ676: *daf-22(m130) II; him-8(e1489) IV; otIs45 [Punc-119::GFP] V*, PB4641: *C. remanei*,

TU3595: *sid-1(pk3321) him-5(e1490) V; lin-15B(n744) X; uIs72 [pCFJ90(Pmyo-2::mCherry) + Punc-119::sid-1 + Pmec-18::mec-18::GFP]*,

CQ697: *daf-22(m130) II; him-8(e1489) IV; vIs69 [pCFJ90 (Pmyo-2::mCherry + Punc-119::sid-1)] V*,

CQ743: *kyIs140 [str-2::GFP + lin-15(+)] I; daf-22(m130) II; him-8(e1489) IV*.

IQ7701: *Phrg-7::GFP*

All strains are maintained using standard methods. C. elegans are maintained at 20℃ on high growth media (HGM, 3 g/L NaCl, 20 g/L Bacto-peptone, 30 g/L Bacto-agar in distilled water, with 4 mL/L cholesterol (5 mg/mL in ethanol), 1 mL/L 1M CaCl2, 1 mL/L 1M MgSO4, and 25 mL/L 1M potassium phosphate buffer (pH 6.0) added to molten agar after autoclaving) or normal growth media (NGM, HGM recipe modified as follows: 2.5 g/L Bacto-peptone, 17 g/L Bacto-agar, and 1 mL/L cholesterol (5 mg/mL in ethanol); all other components same as HGM) seeded with OP50 bacteria (CGC). To synchronize worms, gravid adult Day 2 worms are treated with 15% sodium hypochlorite solution (80mL ddH2O, 15mL NaOCl, 5mL KOH) for 8 minutes, or until 80% of worm body are invisible, to obtain synchronized eggs and avoid overbleaching. Synchronized eggs are directly put on seeded HGM or NGM plates and allowed to grow. For RNA interference assays, worms are grown on HGM plates until L4, and then transferred onto HGM + IPTG+ Carbenicilin plates seeded with respective HT115 RNAi bacteria from the Ahringer Library. For experiments with aged worms, worms are transferred onto plates supplemented with Floxuridine (FUDR) starting from L4. Worms are transferred to plates without FUDR at least 1 day before behavioral experiments. For experiments where a pure male population is needed, mixed Day 1 adult male and hermaphrodites are washed off plates with M9 and additionally washed 2 times before freely passing through a 35mm nylon mesh filter for 10 mins to separate males from hermaphrodites. Males and hermaphrodites are then examined under a microscope for filtration success. A 95% male purity is needed for further experiments.

### Chemotaxis and learning and memory assays

The learning and memory assays were performed as previously described (Kauffman et al., 2010, 2011; Stein and Murphy, 2014; Kaletsky et al., 2016; Weng et al., 2023; Zhou et al., 2023). Briefly, chemotaxis assays towards 10% butanone were performed for both naïve untrained worms and trained worms left on holding plates for various amounts of time to identify naïve chemotaxis, learning, and memory, respectively. Chemotaxis plates were prepared by placing 1uL of 10% butanone diluted in ethanol and 1uL of 100% ethanol on two spots 80mm apart on unseeded 100mm NGM plates. Each spot also had 1uL of 75% NaN3 to immobilize the worms. Worms were washed off plates with M9 and washed 2 additional times to remove residual bacteria before being put on chemotaxis plates on a spot 5.5cm away from either the butanone or ethanol spots and given 1 hour to chemotaxis to butanone or ethanol spots. Chemotaxis was measured as below

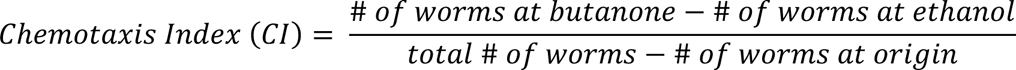

The learning index was calculated as below

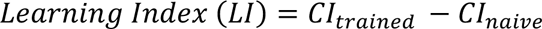

Naïve worms were tested for chemotaxis toward butanone without training. Learning and memory were measured using worms trained on seeded NGM plates with 18 µL butanone streaks on the lid for an hour. Learning was assessed immediately after training, while 1-hour and 2-hour memory were evaluated by placing worms on seeded holding plates without butanone streaks for 1 hour or 2 hours before measuring chemotaxis, respectively.

For learning and memory assays involving male worms, male specimens were filtered a day before the experiment to ensure recovery from filtration.

For pyrazine chemotaxis assays, the procedure was similar to the naïve chemotaxis assay, except 1 µL of 10% pyrazine diluted in ethanol was used instead of 10% butanone.

### Pheromone preference assay

Pheromone preference assays were performed similar to previously described (White et al., 2007; Shi et al., 2017; Wan et al., 2019). For the pheromone preference assay, we used *C. remanei* true female pheromone because it had been shown to be more attractive than hermaphrodite pheromone. *C. remanei* pheromone was collected by separating L4 males or females onto seeded plates without the opposing sex until Day 4. Then Day 4 worms were picked into M9 with a concentration of 10 worms/100uL M9. Worms were kept in M9 overnight to collect pheromone. For pheromone chemotaxis plates, 2uL of pheromone solution and M9 solution were dotted on 2 different spots separated 5cm apart on a 60mm unseeded NGM plate. Each spot also had 1uL of 7.5% NaN_3_ solution to immobilize worms once they reached the spot. Chemotaxis assays to pheromone were performed similarly to chemical chemotaxis assays. Males are washed onto chemotaxis plates and given 1hr to chemotaxis to either spot. The Chemotaxis Index was measured as chemical chemotaxis assays.

### Neuron morphology imaging

We transferred synchronized worms to FUDR at L4 and maintained them by transferring them to fresh plates every 2 days. We imaged worms on adult Day 2 and Day 12, using Nikon AXR confocal microscope’s GFP channel at 60X magnification and 0.5um Z-stack. Images were processed using NIS-Elements and FIJI software and blindly quantified. Beading was measured as bead-like varicosities in the neurite, and wavy neurites were measured as significant path changes from normal neurite processes. Imaging was performed in 3 biological replicates, and 40 Day 2 and 56 Day 12 worms were imaged. Data analysis was performed using Prism software, and Chi-square statistical measurements were used.

### Neuron sorting and RNA extraction

Neuron sorting and RNA extraction were performed as previously described (Kaletsky et al., 2016, 2018). For each biological replicate, 8 synchronized HGM plates of male worms were filtered to obtain a >95% pure male population on Day 1, and neuron sorting was performed on Day 2 and Day 8. Briefly, SDS and DTT-containing lysis buffer was used to break the cuticle, and then 20mg/mL pronase solution from Streptomyces griseus accompanied by mechanical disruption was used to isolate single cells from the tissue. Cells are suspended in L-15 butter with 2% Fetal Bovine Serum (FBS) and kept at 4C throughout the sorting procedure. Cells are sorted using a FACSVantage SE w/ DiVa (BD Biosciences; 488nm excitation, 530/30nm bandpass filter for GFP detection). We determined sorting gates by comparing with age-matched, genotype-matched non-fluorescent samples. The sorting process lasts about 40 minutes. Fluorescent neuron cells were directly sorted into Trizol. 100,000 GFP+ cells were collected for each sample. We collected six biological replicates for each age and sex. We used standard trizol-chloroform-isopropanol procedure for RNA extraction, and RNeasy MinElute Cleanup Kit (Qiagen) for RNA cleanup. RNA quality and concentration were then measured by Qubit and Agilent Bioanalyzer RNA Pico chip.

### RNA sequencing and data analysis

Library preparation was performed using Ovation SoLo RNA-Seq library preparation kit with AnyDeplete Probe Mix-*C. elegans* (Tecan Genomics) with manufacturer’s instructions and 2ng of RNA input was used. Library quality and concentration were assessed using an Agilent Bioanalyzer DNA 12000 chip. Then we multiplexed the samples and performed sequencing using the NovaSeq S1 100nt Flowcell v1.5 (Illumina).

Sequencing data analysis was conducted using the Galaxy website. FastQC was applied to each sample for quality control analysis. The RNA STAR package mapped reads to the *C. elegans* genome (ce11, UCSC Feb 2013) using the gene model ws245genes.gtf, resulting in 50-70% of reads being uniquely mapped. Reads that uniquely mapped to the genome were counted using the htseq-count algorithm in union mode. DESeq2 analysis was then employed for read normalization and differential expression analysis of the counted reads (Love et al., 2014). PCA analysis was performed along with the DESeq2 package. Genes with a log_2_(fold-change) > 1 and p-adjusted <0.05 were considered differentially expressed in subsequent analysis. Gene ontology analysis was performed using either gprofiler (Raudvere et al., 2019) or WormCat 2.0 (Holdorf et al., 2020) category 2. Tissue queries were performed on the top 500 highest fold-change genes, using the worm tissue query website (www.worm.princeton.edu) (Kaletsky et al., 2018), and only major systems were selected in the analysis. The location, transcript and gene length of each gene is obtained from the Wormbase using the Wormmine. Transcript per million is calculated as below

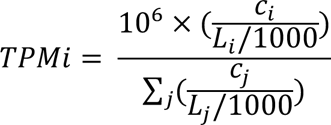

Where TPMi is the Transcripts Per Million for the i-th gene, Ci is the read count for the i-th gene, and Li is the length of the i-th gene in base pairs.

### Quantitative and Statistical Analysis

Experimental analyses were analyzed using the Prism 10 software. Two-way ANOVA with Tukey post-hoc tests were used to compare the learning curve or pheromone preference curve between multiple conditions. One-way ANOVA followed by Dunnet post-hoc tests for multiple comparisons was performed to compare chemotaxis index between control and experimental conditions. Chi-square test was performed to compare the neuron morphology change between young and aged AWC neurons. All GO term analysis were perform using gprofiler with adjusted p-values or Wormcat 2.0 software with Bonferroni corrected adjusted p-values as noted in the figure legend. Venn diagram overlaps were compared using the hypergeometric test. Differential expression analysis of RNA-seq and PCA were performed using the DESeq2 algorithm and adjusted p-values were generated with the Wald test using the Benjamini and Hochberg method (BH-adjusted p-values). Additional statistical details of experiments, including sample size (with n representing the number of chemotaxis assays performed for behavior, sample collections for RNA-seq, and the number of worms for microscopy), can be found in the methods and figure legends. Linear regression, Pearson’s sorrelations and skewness were calculated using the python sklearn and SciPy packages. Skewness is calculated from the distribution histogram of hermaphrodite log(TPM+1) – male log(TPM+1) values.

## Data availability

Sequencing reads are deposited at NCBI BioProject under accession number PRJNA1054152.

## Code availability

Analysis codes generated from this study are available upon request.

## Resource availability

Further information and requests for resources and reagents should be directed to and will be fulfilled by Coleen T. Murphy.

## Supplemental Figures

**Figure S1.**
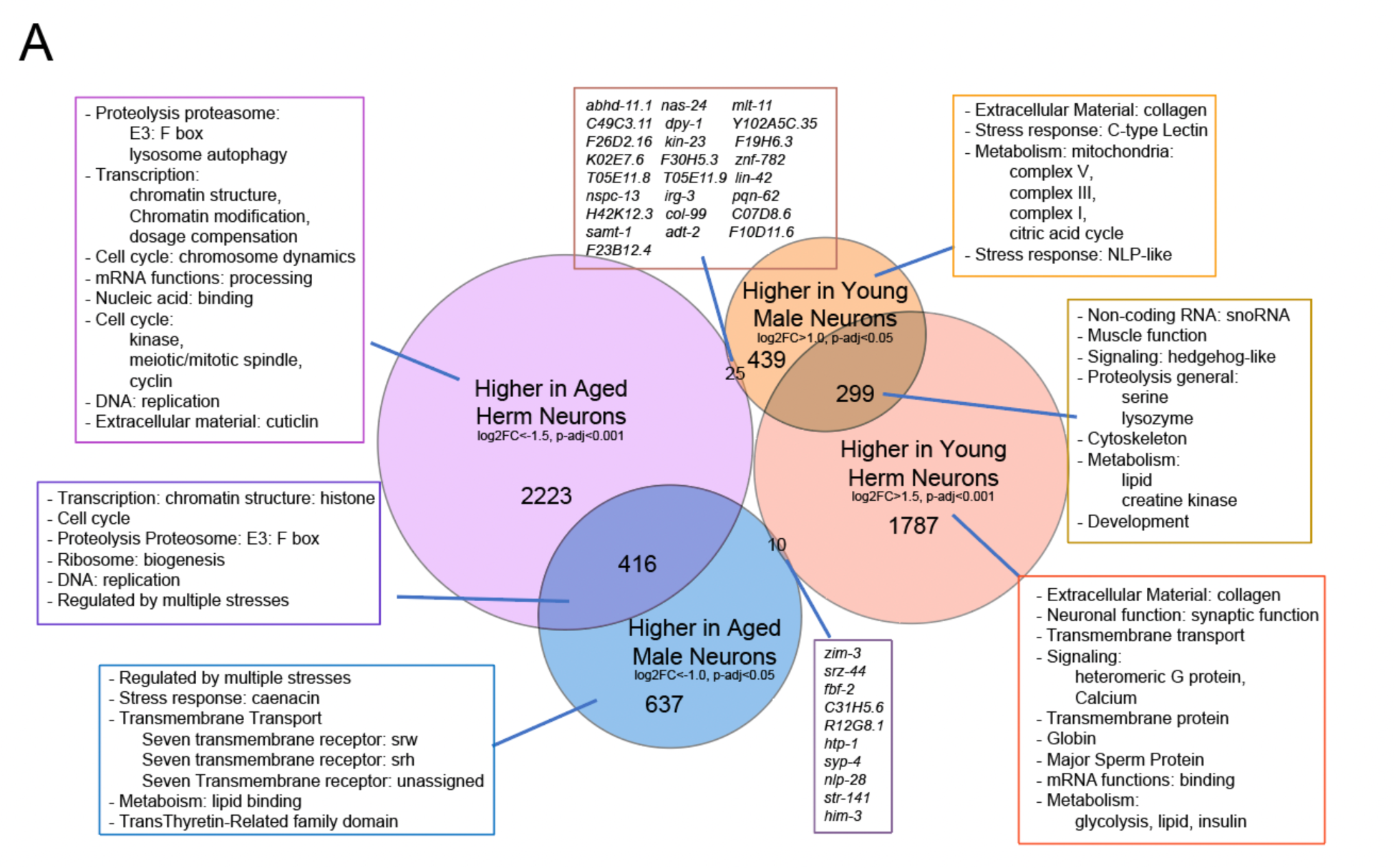
Venn diagram showing overlaps of differentially expressed genes and their gene ontology terms in both sexes. Gene ontology from WormCat 2.0 category 2. p-values of overlapping regions are obtained from the hypergeometric test. Related to Figure 3D.

**Figure S2.**
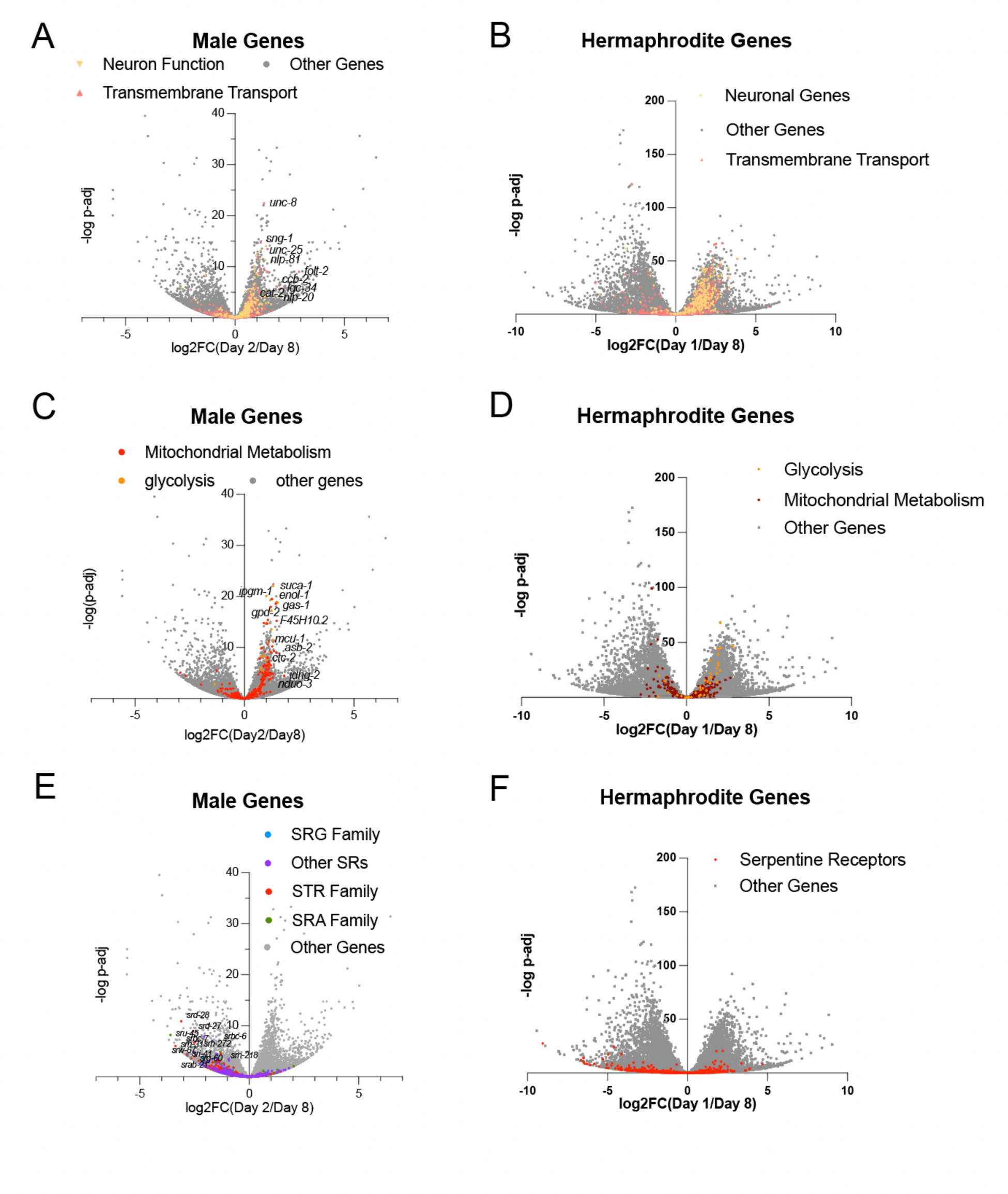
Neuronal genes in enriched GO categories are differentially expressed with age in males. Related to Figure 4 and Figure 5. **(A)** Volcano plot of neuron functional genes and transmembrane transport genes’ expression change with age compared with all genes in males. Related to Figure 4A. (B) Volcano plot of neuron functional genes and transmembrane transport genes’ expression change with age compared with all genes in Hermaphrodites. (C) Volcano plot of serpentine receptor GPCR genes’ expression changes with age compared with all genes in males. Related to Figure 4C. (D) Volcano plot of serpentine receptor GPCR genes’ expression changes with age compared to all genes in hermaphrodites. (E) Volcano plot of mitochondrial metabolic and glycolysis genes’ expression changes with age in males compared with all genes. Related to Figure 5A. (F) Volcano plot of mitochondrial metabolic and glycolysis genes’ expression changes with age in hermaphrodites compared with all genes.

**Figure S3.**
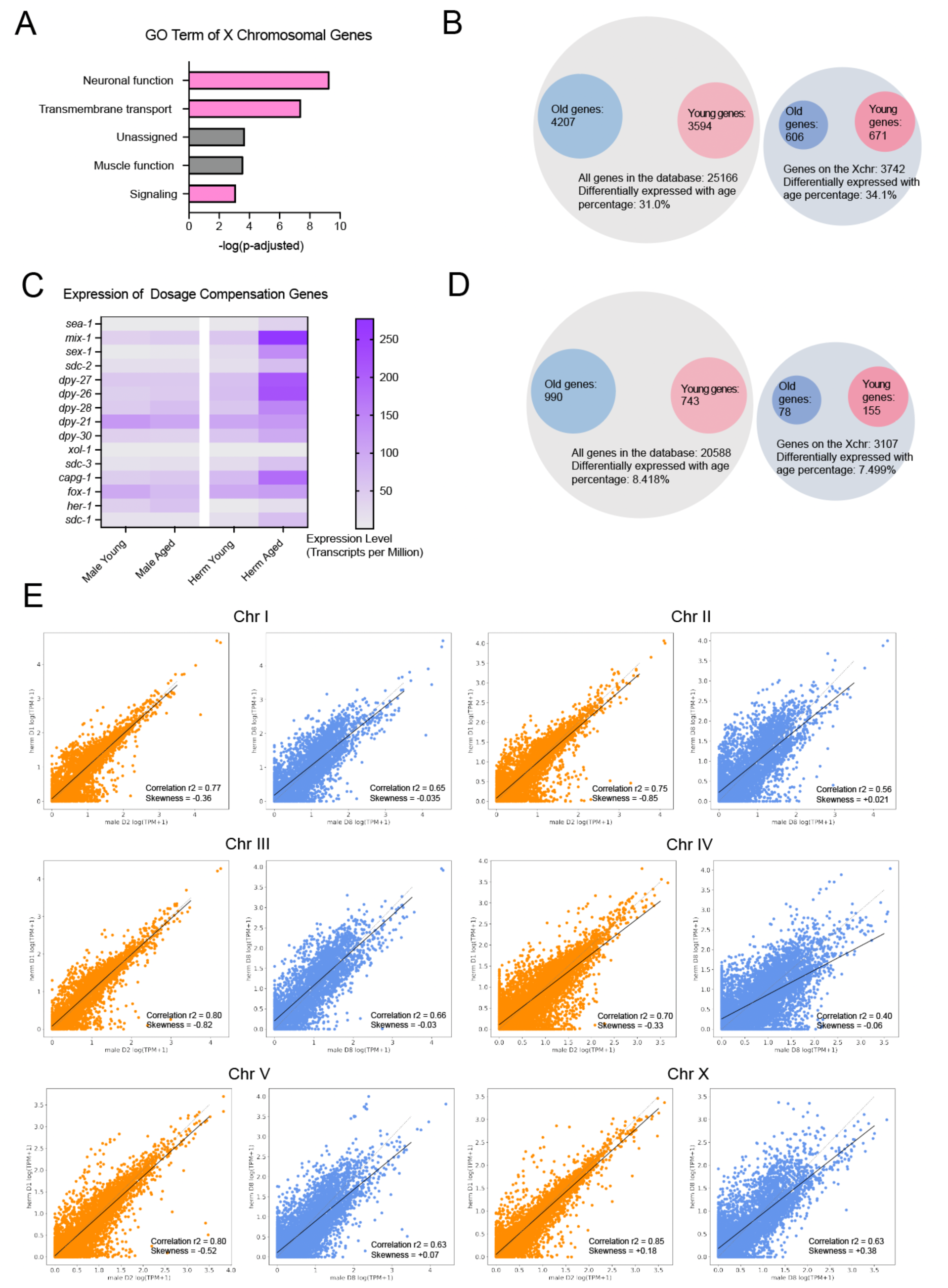
Sex-specific expression changes with age exhibit different characteristics in the X chromosome compared to the autosomes. Related to Figure 6. (A) GO Term of genes on the X chromosome. GO terms are obtained from Wormcat 2.0, category 1. (B) Number of genes differentially expressed in the genome and on the X chromosome in hermaphrodite neurons. (C) Expression of dosage compensation genes from young and aged neurons in both sexes. Expression level is normalized and shown as transcripts per million. (D) Number of genes differentially expressed in the genome and on the X chromosome in male neurons. (E) The correlation of gene expression level between hermaphrodites and males in young and aged animals. Gene expression levels are shown as transcripts per million. Correlation coefficient and skewness are measured as shown in the methods.

**Figure S4.**
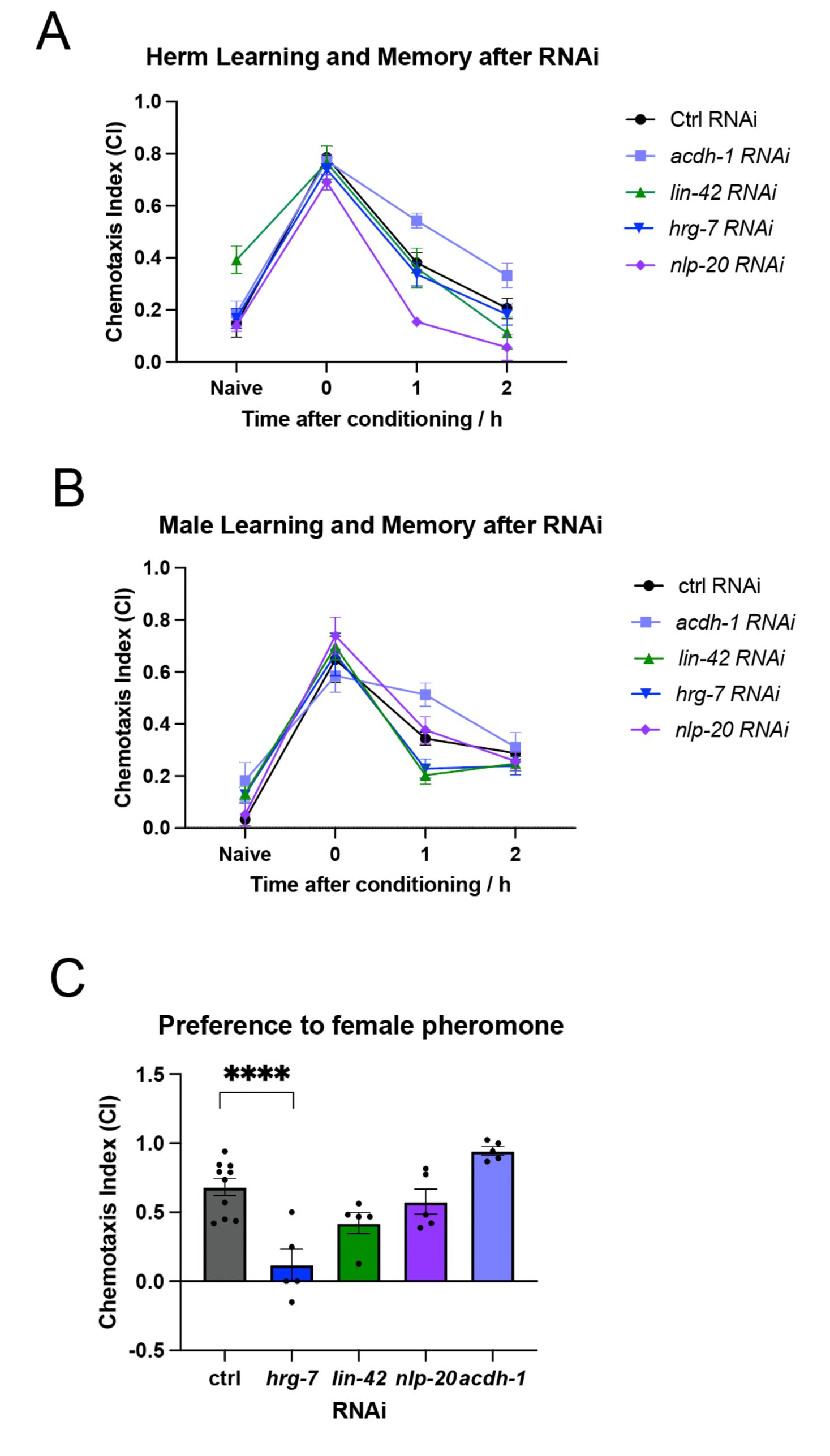
Behavioral test of genes whose expression is higher in young neurons and decreases with age in males. (A) Learning and memory ability of hermaphrodite worms before and after RNAi. (B) Learning and memory ability of hermaphrodite worms before and after RNAi. (C) Pheromone chemotaxis ability of males before and after RNAi of selected genes.

